# miR-21: A therapeutic target for delaying severe liver disease and hepatocellular carcinoma in high-fat-diet-fed mice

**DOI:** 10.1101/2024.09.19.613915

**Authors:** Urmila Jagtap, Anan Quan, Yuho Ono, Jonathan Lee, Kylie A. Shen, Sergei Manakov, Gyongyi Szabo, Imad Nasser, Frank J. Slack

## Abstract

Liver disease, including hepatocellular carcinoma (HCC), is a major global health concern, claiming approximately 2 million lives worldwide annually, yet curative treatments remain elusive. In this study, we aimed to investigate the role of microRNA-21-5p (miR-21) in metabolic dysfunction-associated steatotic liver disease (previously NAFLD), metabolic-associated steatohepatitis (previously NASH), and HCC within the context of a Western high-fat diet, without additional choline (HFD) and offering potential therapeutic insights. We found that reduced miR-21 levels correlated with liver disease progression in WT mice fed on HFD, while miR-21 knockout mice showed exacerbated metabolic dysfunction, including obesity, hepatomegaly, hyperglycemia, insulin resistance, steatosis, fibrosis, and HCC. Our study reveals that miR-21 plays a protective role in metabolic syndrome and in the progression of liver disease to cancer. MiR-21 directly targets *Transforming growth factor beta-induced* (*Tgfbi*), a gene also known to be significantly upregulated and a potential oncogene in HCC. Further, our study showed that intervention with the administration of a miR-21 mimic in WT livers effectively improves insulin sensitivity, steatosis, fibrosis, *Tgfbi* expression and tumor burden in HFD conditions. These findings indicate that miR-21 could serve as an effective strategy to delay or prevent liver disease in high-fat-diet environments.

**Significance:** Our study demonstrates in vivo that miR-21 has protective functions in the broad spectrum of high-fat diet-based, progressive liver disease and cancer, and we show potential therapeutic value of a microRNA-21 mimic.

**Graphical abstract:** 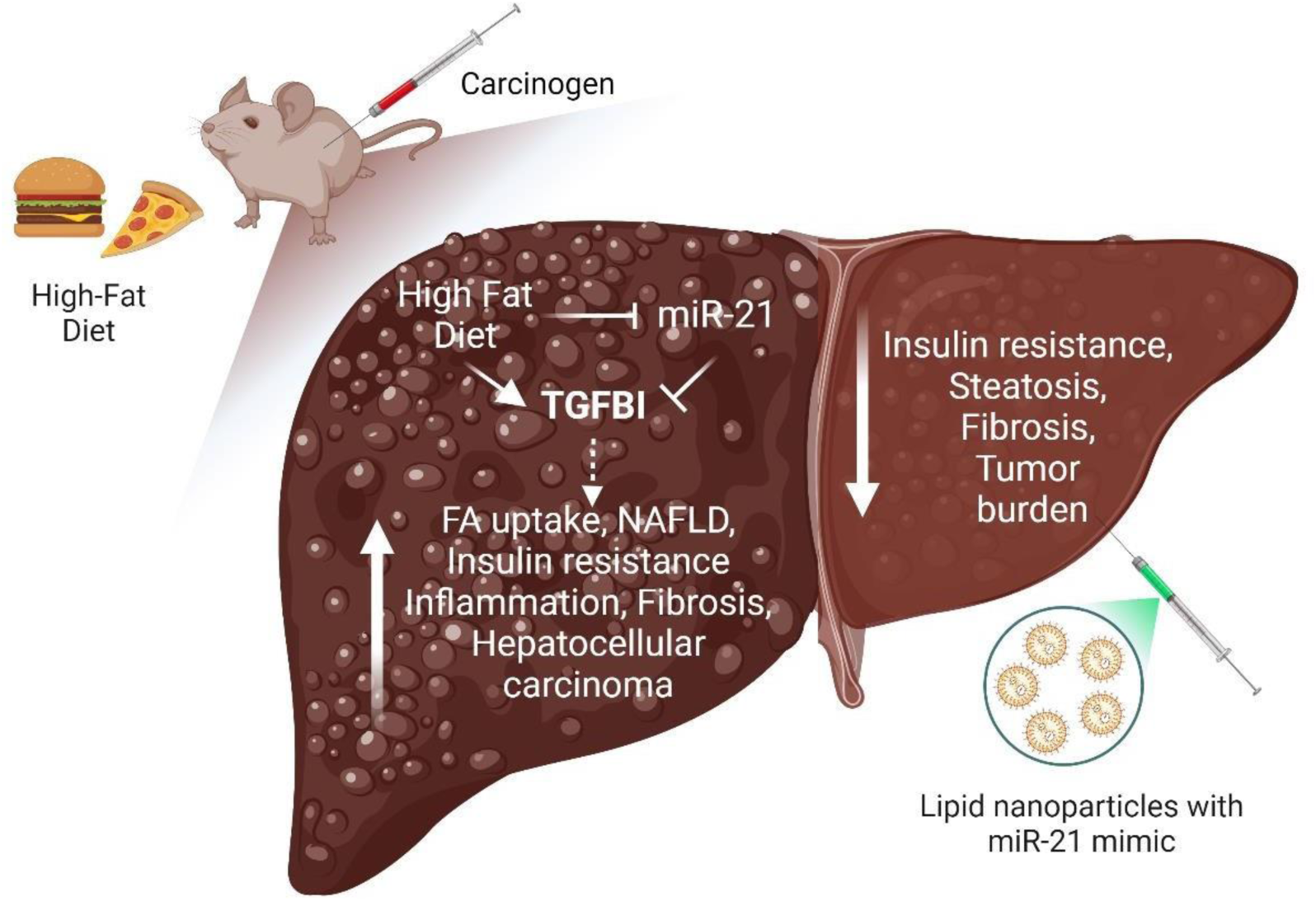

## Introduction

Liver disease is one of the greatest contributors to the global health burden (1) and hepatocellular carcinoma (HCC) is the 4^th^ most prevalent cancer (2). Liver disease manifests as metabolic dysfunction-associated steatotic liver disease (MASLD) (formerly, non-alcoholic fatty liver disease-NAFLD) or Met-ALD, MASLD with alcohol (formerly, alcoholic steatohepatitis-ASH) (3) and pathophysiology includes a gradual progression from simple fatty liver, i.e., steatosis, to steatohepatitis, fibrosis, cirrhosis, and terminally, to hepatocellular carcinoma (4,5). In addition to genetic predisposition, hepatic insults in the form of a fatty diet, alcohol, drug intake, and viral infections are largely responsible for hepatic dysfunctions (6). In addition to affecting liver pathophysiology, these factors are also correlated with rising rates of metabolic disorders (7), like obesity, hypertension, hyperglycemia, hyperlipidemia, and insulin resistance (8), together resulting in significant disease morbidity and mortality. Indeed, the Western diet consisting of high caloric value has claimed more than 100 million individuals in the United States alone (9).

It is well-documented that consumption of a high-fat diet (or so-called Western diet) leads to liver damage (10,11) and can be described by a multiple-hit model (12). The primary insult originating from genetic predisposition sensitizes the liver to injury. The secondary hits by subsequent insults like inflammatory cytokines, adipokines, endoplasmic reticulum stress, oxidative stress, mitochondrial dysfunction, and hepatocellular death then lead to impaired fatty acid metabolism, and insulin resistance triggering steatosis, steatohepatitis, and fibrosis (11).

MicroRNAs are a class of small non-coding RNA molecules that play various roles in metabolic functions. One such microRNA is microRNA-21 (miR-21), which has been shown to function in glucose metabolism, fatty acid uptake, metabolism, inflammation, etc. (13,14). The dysregulation of miR-21 promotes insulin resistance, hepatitis, and fibrosis in various cell lines and mouse models (15). In addition, increased hepatic expression in animal models and patients with MASLD/MASH and HCC has been reported (13,14); however, the serum levels of miR-21 vary significantly in different studies (15). Contradictory reports exist correlating liver disease with both increased (16,17) and decreased (18) expression of miR-21 in human serum. However, despite the possible correlation of miR-21 levels with liver disease, efforts to utilize it as a diagnostic marker or therapeutic intervention for liver disease and HCC have so far failed (16,18,19). One of the most likely reasons is the high variability of miR-21 in the serum and tissue of patient samples. Studies over more than a decade have hinted towards the possible context-dependent functions of miR-21 in various liver diseases like HCC, probably based on the nature of the insult (20–22).

The function of miR-21 as an oncogene and hepatotoxic agent is a long-standing dogma. It was proposed that miR-21-5p promotes NASH-related hepatocarcinogenesis via PPARα (23) or through miR-21 interaction with the Hbp1-p53-Srebp1c pathway, highlighting that miR-21-ASO could be of potential therapeutic value (24), or that miR-21 promotes migration and invasion in HCC through the miR-21-PDCD4-AP-1 feedback loop (25). At the same time, several reports have highlighted that liver miR-21 progressively increases with NAFLD/MASLD progression, acting upon specific pathogenic pathways at each disease stage (26). For example, it has been shown that PPARɣ attenuates inflammation and oxidative stress in NASH by inhibiting the miR-21-5p/SFRP5 pathway (27), that miR-21 abrogation, decreasing PPARα, significantly improves whole-body metabolic parameters in NASH (28), that miR-21 closely associates with fibrosis in a rat model of NASH (29), hepatic miR-21/miR-21* deficiency prevents glucose intolerance and steatosis in mice fed on obesogenic diet (30), and inhibiting or suppressing liver miR-21 expression reduced liver cell injury, inflammation and fibrogenesis in distinct NASH models by targeting PPARα expression (31).

However, recently emerging reports have challenged this belief, while some show no dependency on miR-21 on liver fibrosis and other phenotypes (32). Contradictory to reports correlating liver disease with increased levels of miR-21 in patient serum samples, Chuanzheng Sun et al. report reduced levels of miR-21 in serums of NAFLD patients compared with the healthy control. Furthermore, they demonstrate that miR-21 negatively regulates the levels of triglycerides, free cholesterol, and total cholesterol in PA/OA-treated HepG2 by attenuating its direct target HMGCR (33).

Recently, two independent reports showed results in contrast to the expected oncogenic role of miR-21. Using two different models of hepatocellular carcinoma, i.e. genetic and carcinogen (Diethylnitrosamine, DEN) based, Mart Correia de Sousa et al. showed that total or hepatocyte-specific genetic depletion of miR-21 fosters HCC development (34). Another study by Said Lhamyani et al. demonstrated that miR-21 activates a thermogenic program in brown adipose tissue and browning in different white adipose tissue depots, suggesting an increase in adipose energy expenditure (35). These studies reveal the importance of *in vivo* experiments in revealing functions for miR-21. Until our study, the role of miR-21 in liver disease and cancer under prevailing human physiological western diet conditions has not been examined.

Considering the high prevalence of liver disease and HCC worldwide and the lack of preclinical alternatives, animal models that can recapitulate the pathophysiology of this progression are essential. In this study, we have created a mouse model of liver disease and cancer that shows all classical metabolic and pathological features associated with various stages of human liver disease. Over the course of 32 weeks following a one-time DEN administration and a high-fat-diet-based protocol, mice sequentially developed metabolic phenotypes starting from obesity, hyperglycemia, and insulin resistance to the entire spectrum of liver disease, i.e., steatosis, fibrosis, MASH, and ultimately, hepatocellular carcinoma. Using bulk-RNA sequencing of the liver representing various stages of liver disease, we propose pathways that could contribute to transitioning the healthy liver to the progressive disease condition upon HFD insult. Since we found that miR-21 levels decrease over time in this model, we investigated the role of miR-21 on the progression of liver disease using this protocol with a whole-body knockout of miR-21.

Our results indicate that the loss of miR-21 is pathogenic to the liver and causes metabolic dysfunction in mice. The severity of liver disease increased significantly when the mice were fed a high-fat diet, representing multiple insults to the liver and resulting in a more severe diseased condition. In our transcriptomics analysis, we identified the molecular pathways regulated by miR-21 and HFD independently and in combination. Through various analyses, we propose that one key mechanism of miR-21 action is via directly targeting the *Transforming growth factor beta-induced* (*Tgfbi*) transcript. Further, we investigated whether administration of miR-21 in the form of mimic in the WT mice can protect from or delay liver disease pathogenesis and HCC in HFD. We saw increased insulin sensitivity, reduced steatotic and fibrotic phenotypes, and reduced tumor burden in these livers. Together, our study has revealed a protective role of miR-21 in liver disease and HCC in the HFD setting and advocates it as a target to delay liver disease progression.

## Results

### WT mice on HFD develop obesity, liver injury, hyperglycemia, and insulin resistance

Generating an accurate model of human liver disease progression to HCC was one of the aims of this study. B6/129S (WT) male mice were given a one-time dose of 25mg/kg of Diethylnitrosamine (DEN) intraperitoneally at three weeks of age. Following injections, all mice were randomly divided into two feeding categories – chow or high-fat diet, without additional choline (HFD) (Fig. 1A). All the mice were weighed weekly, and the body weight was graphed over time. As expected, the HFD-fed mice showed significantly higher weights (Fig. 1B) as well as increased body-fat accumulation (Suppl. Fig. 1E) over time than chow-fed mice, indicating obesity in this group. The obesity phenotype is usually associated with hepatomegaly and liver injury. Interestingly, the liver-to-body mass ratio did not showany increase upon HFD feeding, indicating that these mice did not develop hepatomegaly (Fig. 1C). However, the measurement of ALT levels in serum showed a significant increase, highlighting liver injury (Fig. 1D). Metabolic dysfunctions that are known to be associated with high-fat diet consumption were also tested. The increased concentrations of blood glucose indicated that the mice fed on a high-fat diet suffered from hyperglycemia compared to chow-fed counterparts (Fig. 1E). The insulin tolerance test representing insulin resistance (Fig. 1F) showed delayed insulin response in mice fed with HFD.

**Figure 1:**
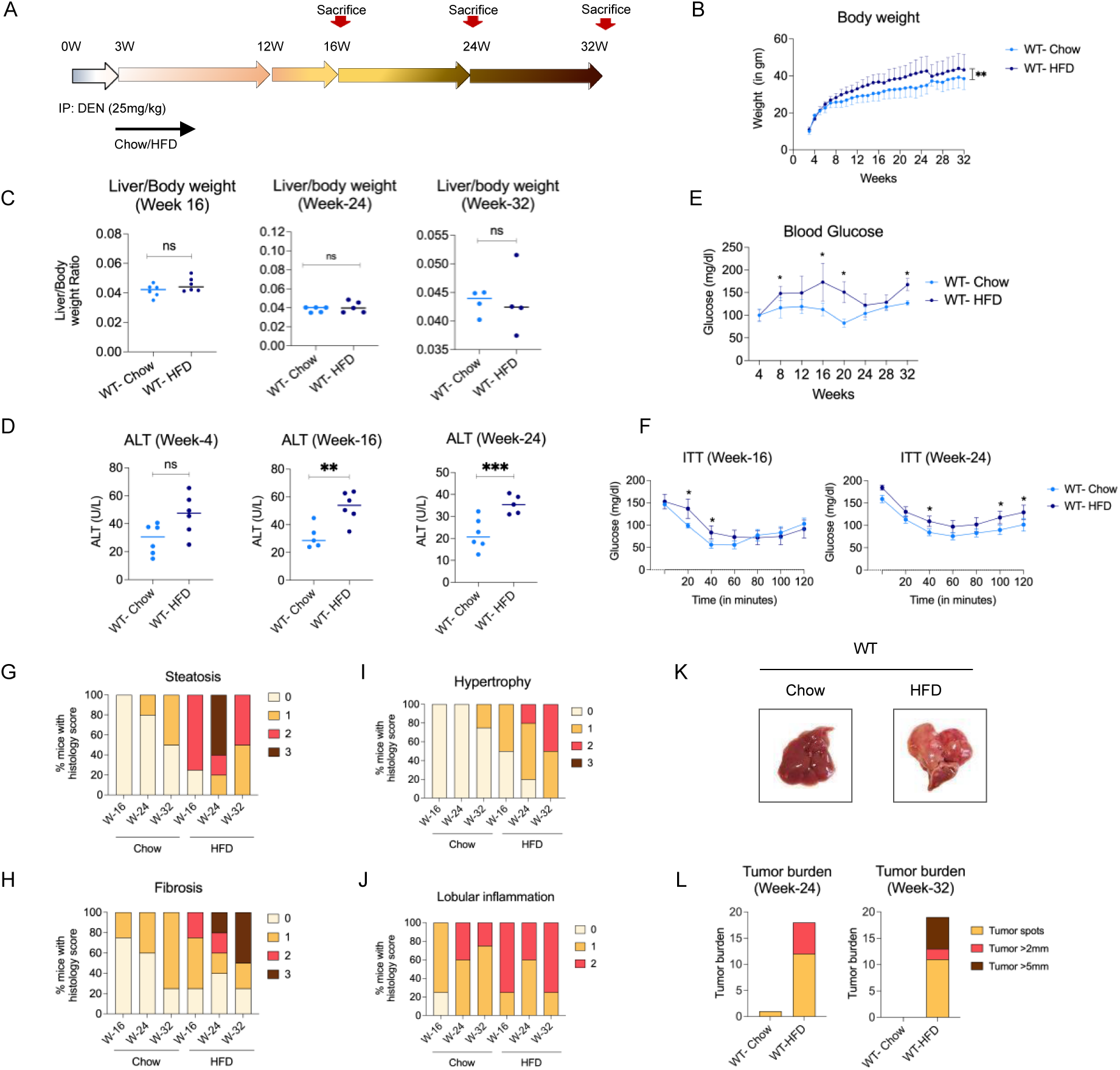
WT mice fed on HFD develop obesity, liver injury, hyperglycemia, insulin resistance, fatty liver, steatohepatitis, advanced fibrosis, and HCC. (A) Experimental timeline. B6/129S mice were injected with 25mg/kg of Diethylnitrosamine (DEN) at three weeks of age and were fed with either a chow diet (Chow) or high-fat diet, without additional choline (HFD) for up to 32 weeks, (B) change in body weight over time, (C) Liver to body weight ratio, (D) Serum ALT levels,(E) Blood glucose levels, and (F) Insulin tolerance test (ITT) for week-16 and week-24. Histology scores for (G) Steatosis,(H) Fibrosis, (I) Hypertrophy, and (J) Lobular inflammation. (K) Representative liver images at week 32, and (L) Tumor burden at weeks 24 and 32. Data are expressed as the mean±SD for six mice per group for ITT assay, at least six mice per group for body weight measurements, histology and tumor burden analysis. \**p*<0.05, \*\**p*<0.01, \*\*\**p*<0.001.

### WT mice on HFD sequentially develop steatosis, steatohepatitis, advanced fibrosis, and liver tumors

A set of mice were euthanized at each time point, i.e., week 16, 24, and 32, and the livers were analyzed using H&E, Oil Red O, and Sirius red stains to assess liver histopathology, inflammation, fat accumulation, and fibrosis, respectively. Histopathological analysis indicated that starting from week 16, as compared to chow-fed counterparts, all WT mice fed on HFD showed the presence of mixed micro-vesicular and macro-vesicular fat droplets within hepatocytes throughout the liver, as also confirmed by Oil Red O staining (Fig. 3A, 1G). The severity of this phenotype increased, as indicated by the accumulation of stage 3 phenotypes at week 24 (Fig. 3A, 1G). As assessed by Sirius red staining, the presence of stage 1 and 2 fibrosis was apparent in week 16 (Fig. 3A,1H), indicating that the fibrotic phenotype was initiated before week 16. Notably, high scoring of hypertrophy (Fig. 1I) and lobular inflammation phenotypes (Fig. 1J) were also observed in these mice, consistent with MASH. Along with MASH and increased fibrosis in these livers, the HFD-fed mouse livers developed tumor nodules by week 24 (Fig. 1K), which were virtually absent in the Chow group (Fig. 1L, left graph). By week 32 (Fig. 1L, right graph), the tumors progressed to adenomas/dysplastic nodules (∼40%) and fatty adenomas (∼45%). Some of the tumors also showed further advanced progression to the early HCC and/or HCC (∼15%).

### WT mice on HFD show gene expression changes related to metabolic response early in hepatic injury

To better understand the pathways and factors involved in liver disease progression, we isolated RNA from livers from each sub-group at each stage of the protocol and performed RNA sequencing (RNAseq). RNAseq analysis of liver samples showed a significant number of genes to be differentially regulated at each stage (Suppl Fig. 2, Suppl. Fig. 3A). IPA analysis of these genes showed enrichment of pathways involved in fatty acid synthesis and metabolism (Suppl. Fig. 3B, 3D), and inflammation (Suppl. Fig. 3B, 3C) correlating with the steatosis and MASH phenotype observed in the tissue sections in WT-HFD group as compared to the WT-Chow group, especially at week-16, the first time point tested. Interestingly, the greatest metabolic differences between Chow-fed and HFD-fed animals were observed in the week-16 livers. However, the extent of the metabolic differences seems to decrease at weeks 24 and 32.

### Knockout of miR-21 causes metabolic dysfunction and progressive liver disease from fatty liver to advanced fibrosis

Elevated levels of miR-21 in circulation (16,36) and liver tissue (17,37) have been correlated with liver disease in humans. We, therefore, measured miR-21 levels in the liver tissue of our mice by qRT-PCR. Interestingly, in our model, as liver disease progressed from simple steatosis to hepatocellular carcinoma in response to HFD feeding, i.e., from week 16 to week 32, miR-21 levels showed a progressive decrease in expression to almost undetectable levels in week-32 livers (Fig. 2A). Interestingly, we also found several miR-21 targets to be differentially regulated in these livers. (Suppl. fig. 4, 4B, 4C). The KEGG and GO analysis of these genes showed that they are involved in critical metabolic liver functions like lipid metabolism and peroxisomes (Suppl. fig. 4D, 4E). Based on these results, we investigated the role of miR-21 in liver disease progression and performed similar assays as described in Figures 1 and 2 on mice with whole-body knockout of miR-21 (miR-21 KO). The body weight analysis of miR-21 KO mice fed on the chow diet showed no significant change as compared to the WT-Chow group (Fig. 2B, Light red line, Suppl Fig. 5A), indicating no obvious obesity phenotype.

**Figure 2:**
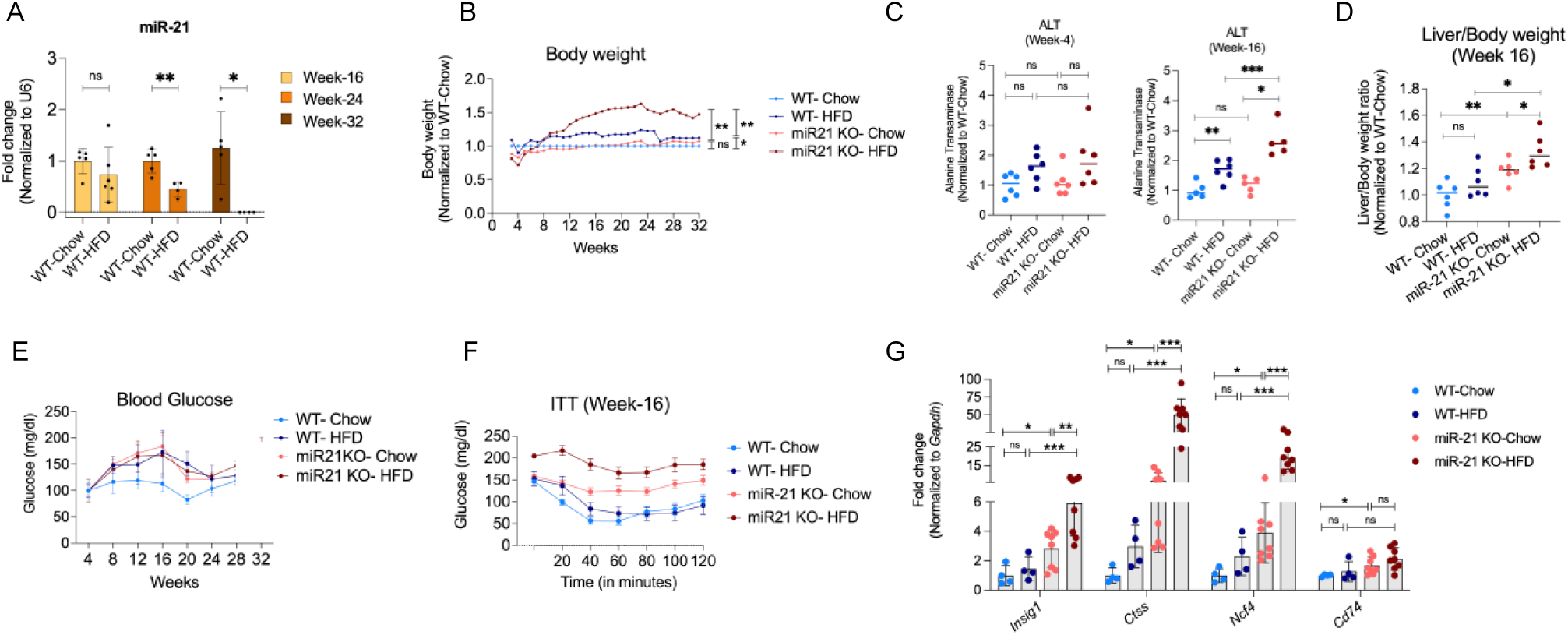
miR-21 KO mice on HFD show exacerbated signs of liver injury, hyperglycemia, and insulin resistance. (A) miR-21 levels in livers during liver disease progression in WT mice (B) Body weight of miR-21 KO mice compared to WT on Chow and HFD group. (C) Serum ALT levels at weeks 4 and 16, (D) Liver to body weight ratio (E) Blood glucose measurements, (F) Insulin tolerance test, and (G) qRT-PCR expression of genes involved in insulin resistance. N= 6-8 mice/group. ns=not significant, *p<0.05, **p<0.01, ***<0.001.

Similarly, no significant changes in serum ALT levels were observed, indicating that these mice had healthy livers at the commencement of the experiment (Fig. 2C). Serum ALT levels showed no change until week 16; however, a significant increase was observed in miR-21-Chow mice as compared to the WT-Chow group at week 24, indicating liver injury. Interestingly, unlike the WT-HFD group, a higher liver-to-body mass ratio was observed in miR-21 KO-Chow-fed mice when compared with the WT counterparts, indicating a hepatomegaly condition in miR-21 KO livers (Fig. 2D). Knockout of miR-21 was also associated with increased blood glucose concentrations, indicating hyperglycemia (Fig. 2E). Insulin tolerance assay showed that miR-21 KO mice were more resistant to the presence of insulin (Fig. 2F). This was also supported by a significant increase in the expression of the genes involved in insulin resistance (Fig. 2G).

H&E staining of miR-21 KO mice fed on the chow diet showed progressive alteration of liver histology as compared to WT counterparts (Fig. 3A). The histological assessment displayed higher scores for hypertrophy (Fig. 3B) and lobular inflammation (Fig. 3C). Increased levels of steatosis and fibrosis were noted in these livers as represented by histo-pathological scoring of H&E/Oil-Red-O (Fig. 3A, 3D, 3F) and Sirius red staining (Fig. 3A, 3E, 3G) as well as qRT-PCR of relevant gene markers (Suppl Fig. 5H, 5I). All these phenotypes increased in severity with time, indicating that loss of miR-21 is sufficient for the gradual progression of liver disease. Though miR-21 KO mice did not show an overt obesity phenotype, the fat mass of these mice was significantly higher than their WT counterparts (Suppl. Fig. 5G). To further evaluate the nature of fat composition in these mice, we measured the serum levels of cholesterol and triglycerides. A significant increase in the serum cholesterol levels was observed in the WT-HFD and miR-21 KO-Chow group as compared to the WT-Chow group (Fig. 3H). Interestingly, however, the triglyceride levels did not differ in any of the groups (Fig. 3I).

**Figure 3:**
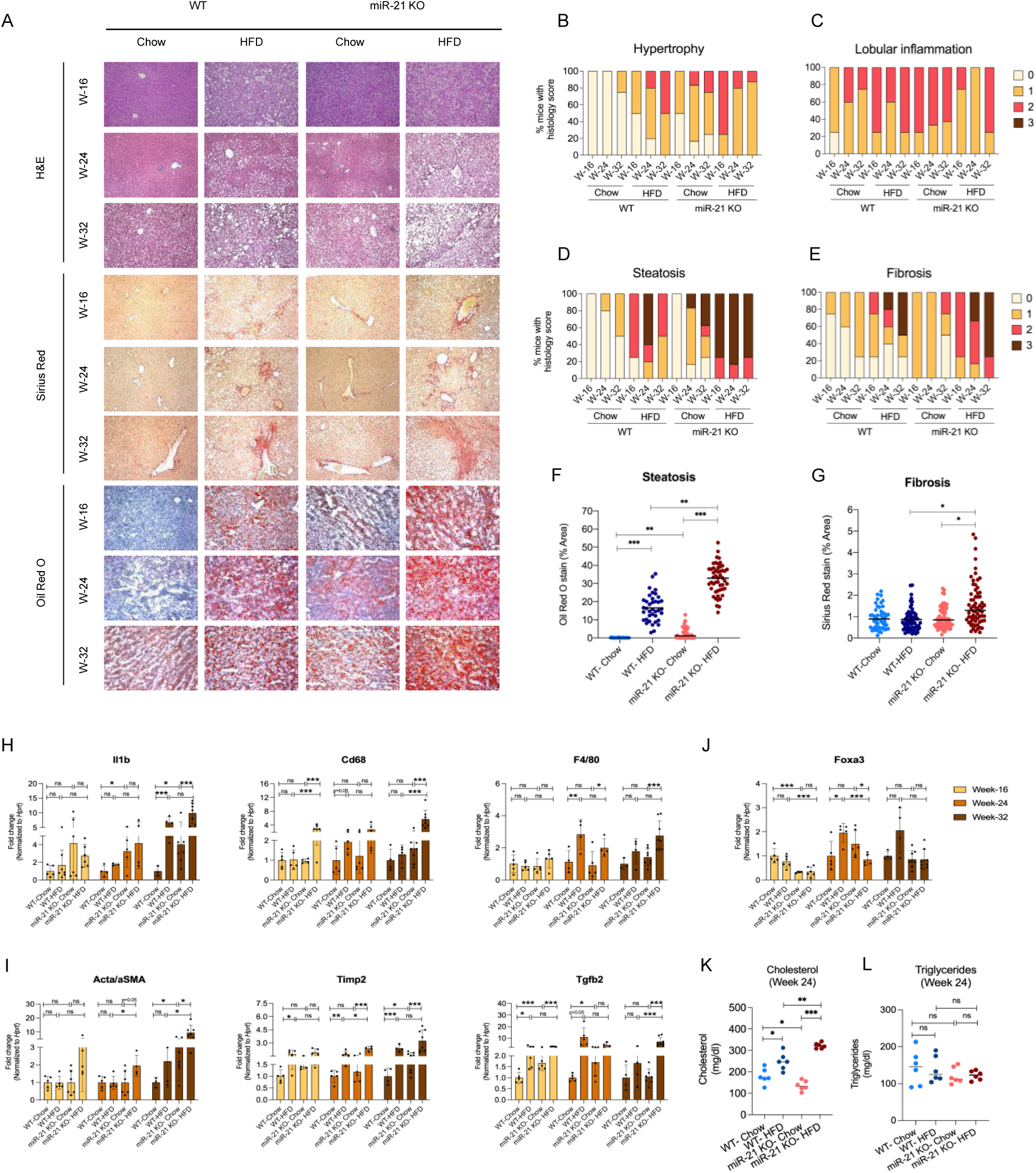
miR-21 KO mice on HFD show exacerbated signs of fatty liver, steatohepatitis, and advanced fibrosis. (A) Representative images of livers at weeks 16, 24, and 32 for Hematoxylin-eosin (H&E), Sirius red, and Oil Red O staining. Histology scores for (B) Hypertrophy, (C) Lobular inflammation, (D) Steatosis, and (E) Fibrosis. ImageJ quantification of (F) Oil Red O and (G) Sirius Red staining at week 16. Quantitative real-time analysis of genes involved in (H) Inflammation (I) Fibrosis, and (J) Fatty acid uptake in Week-16, 24, and 32 liver samples. Serum levels of (K) Cholesterol and (L)Triglycerides at week 24. N= 4-8 mice/group. ns=not significant, *p<0.05, **p<0.01, ***p<0.001.

### miR-21 KO mice on HFD show exacerbated signs of metabolic syndrome, fatty liver, steatohepatitis, advanced fibrosis, and tumor development

We found that the combination of mir-21 KO and HFD showed the highest levels of all the metabolic dysfunction phenotypes we tested. The body weights of miR-21 KO-HFD showed a dramatic increase, with a few of the mice weighing as much as 57gm (Fig. 2B, Suppl. Fig. 5A). Similarly, the measurements of ALT and AST levels in serum showed almost double the levels as compared to the WT equivalents (Fig. 2C, Suppl Fig. 5D, 5E). The blood glucose concentrations, however, did not show an increase over and above the miR-21 KO-Chow group (Fig. 2E). However, the ITT assay showed almost no sensitivity to the presence of insulin at week 16 (Fig. 2F), suggesting that insulin resistance started occurring earlier in the protocol. A qRT-PCR analysis for genes involved in insulin resistance showed high expression (Fig. 2G), corroborating the ITT assay results.

The histological analysis also showed that the miR-21 KO mice fed on HFD have the most severe pathological phenotypes compared to other groups. The H&E, Oil Red O, and Sirius red staining, along with the histological scoring, indicated that the steatotic (Fig. 3A, 3D, 3F) and fibrotic (Fig. 3A, 3E, 3G) phenotypes in miR-21 KO-HFD are initiated earlier and progressed to up to grade 3 severity faster as compared to all other groups. The histology scoring did not show severe lobular inflammation (Fig. 3C), but mRNA quantitative measurements show highly elevated expression of genes involved in inflammation (Suppl Fig. 3H). As described earlier, consistent with their highest fat accumulation, the miR-21 KO-HFD group showed the highest cholesterol levels in serum (Fig. 3H); however, no such increase was observed in the triglyceride levels (Fig. 3I).

### miR-21 KO and HFD have additive effects on the progression of liver disease from steatosis to the development of hepatocellular carcinoma

Throughout all the metabolic and pathological stages of liver disease progression that we tested, the miR-21 KO-HFD group scored the highest in terms of the disease severity as well as the rate of disease progression (Figs. 1 to 3). With regard to tumor burden, we observed drastic differences when comparing the HFD feeding, knockdown of miR-21, and the combination of the two. As mentioned earlier, HFD feeding of WT mice resulted in tumor burden by week 24 (Fig. 1L). Interestingly, in contrast to WT mice, mice from the miR-21 KO-Chow group developed more tumors, indicating the independent role of miR-21 in the progression of liver disease from fibrosis to tumor development (Fig.4A, 4B). As observed in all stages of liver disease, the miR-21 KO-HFD group showed the highest tumor burden compared to any other group. Additionally, the tumors were notably bigger in size (Fig. 4A, 4B). By week 32, some of the tumors reached a diameter of more than 1cm (Fig. 4A). This also explains the lower grading of hypertrophy and lobular inflammation phenotypes in groups fed with HFD, especially miR-21 KO-HFD, where the tissue parenchyma was engulfed in the tumor volume (Fig. 3B, 3C). We did not observe metastasis to lungs or bones in any of the groups.

**Figure 4:**
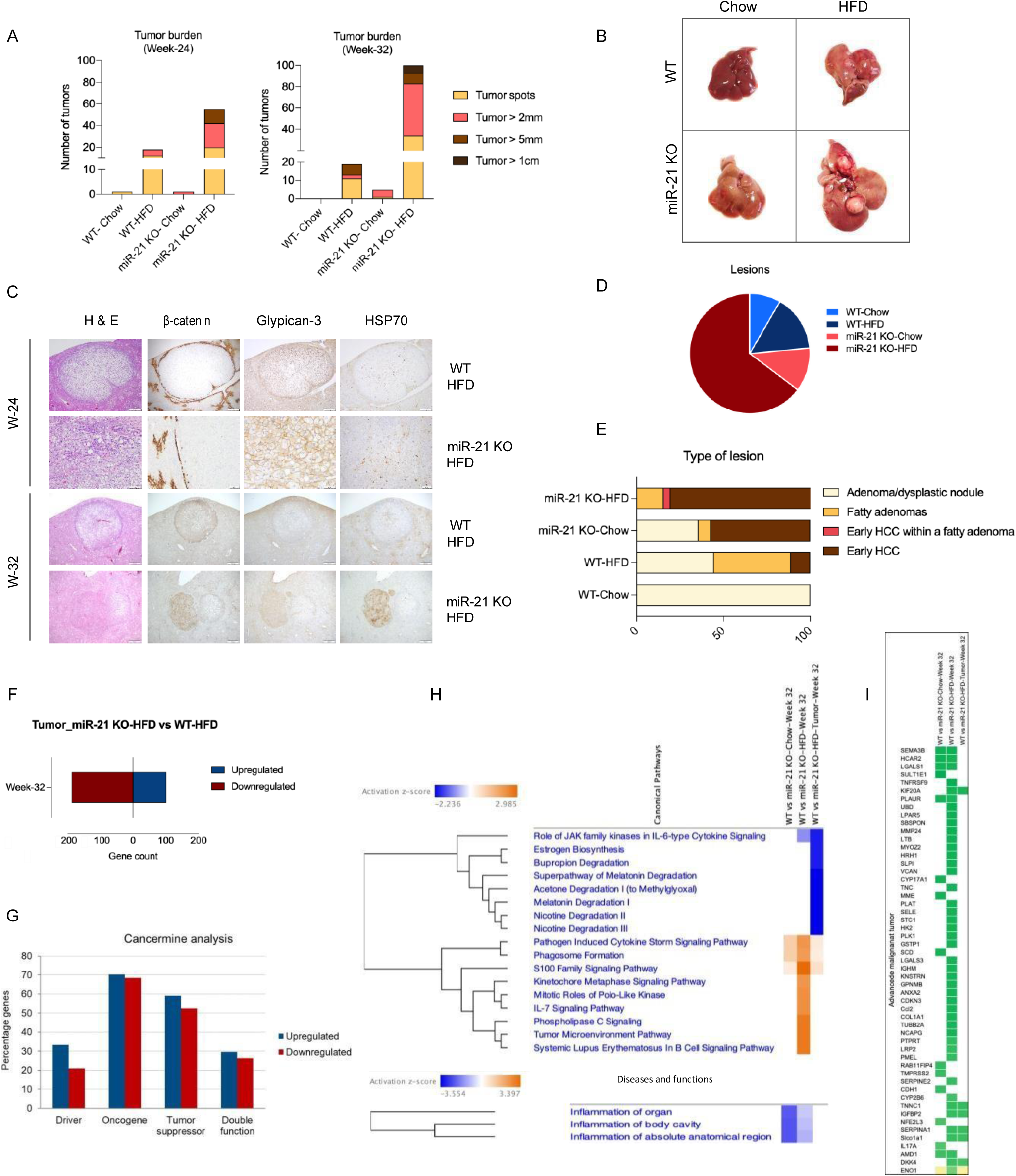
miR-21 KO mice develop hepatocellular carcinoma. (A) Tumor burden of mice at weeks 24 and 32 (B) Representative hepatic tumors (C) H&E and immunostaining of hepatic-tumor sections at weeks 24 and 32, (D) Comparative lesion burden and (E) Types of lesions in various groups, (F) Gene count of differentially expressed genes with fold change >2 in tumors from miR-21 KO vs WT fed on HFD, (G) Cancermine analysis for the nature of genes. (H) IPA analysis of canonical pathways, and diseases and functions, (I) Heatmap representing differentially expressed genes related to the advanced malignant tumor. N= 4-8 mice/group.

Histopathological evaluation of liver tissue sections (Fig. 4C) also corroborated the finding of increased tumor burden, as a greater number of neoplastic lesions were observed in the WT-HFD and miR-21 KO-Chow groups as compared to the WT-Chow group, with the highest being the miR-21 KO-HFD group (Fig. 4D). The tumor profiles of the miR-21 KO-Chow group showed a combination of adenomas/dysplastic nodules (∼36%), fatty adenomas (∼7%), and early hepatocellular carcinomas (∼57%). Interestingly, the tumors were more advanced in this group as compared to the WT-HFD group, with ∼57% early HCC in the miR-21 KO group vs. ∼11% in the WT-HFD group. (Fig. 4E). The combination of the two, however, as in the miR-21 KO-HFD group, showed the most severe pathology, with almost 80% of the tumors already progressed to early hepatocellular carcinoma and only around 20% being fatty adenomas (Fig. 4C, 4E). A small fraction of tumors also showed early HCCs arising inside adenomas (Fig. 4E). All these results indicate that miR-21 and a high-fat diet independently cause pathological effects of tumor development and severe additive effects when combined.

To understand the contribution of miR-21 in causing these tumors, we performed a comparative analysis of our RNAseq data of tumors isolated from miR-21 vs. WT mice fed on HFD. A total of 102 genes were upregulated, and 187 genes were downregulated (Fig. 4F). Cancermine analysis (38) showed 27 up and 19 downregulated genes to have published, supporting literature. Among these upregulated genes, 33% are known to have cancer-promoting driver mutations, 70.3% with oncogenic, 59.2% with tumor suppressor, and 29.6% to have dual reported functions (Fig. 4G, Suppl Table 2). IPA analysis showed significant enrichment in the genes involved in advanced malignant tumors in miR-21 KO liver tumors as compared to WT tumors at week 32. Interestingly, many of these changes are observed in the liver tissue but not inside the tumors (Fig. 4H, 4I), indicating that the liver cells might undergo metabolic changes that drive them to develop tumors.

### miR-21 directly regulates *Tgfbi*

miR-21 KO mice are predisposed to metabolic syndrome and liver disease upon HFD consumption. In order to understand the susceptibility of miR-21 KO mice to liver disease, we compared the gene expression changes in miR-21 KO mice fed on the Chow diet to those of their WT counterparts. As seen previously (Fig. 4), the gene signatures corroborated the disease phenotypes observed in the miR-21 KO mice. Significant upregulation was observed in the genes involved in hepatic steatosis, liver toxicity, and inflammation (Suppl Fig. 5A, 5B, 5C). Additionally, genes involved in retinol biosynthesis, triglyceride degradation, gustation pathway, and protein kinase signaling showed significant upregulation (Suppl Fig. 6), supporting our hypothesis that loss of miR-21 is sufficient to promote the pathogenesis of liver disease.

**Fig. 5:**
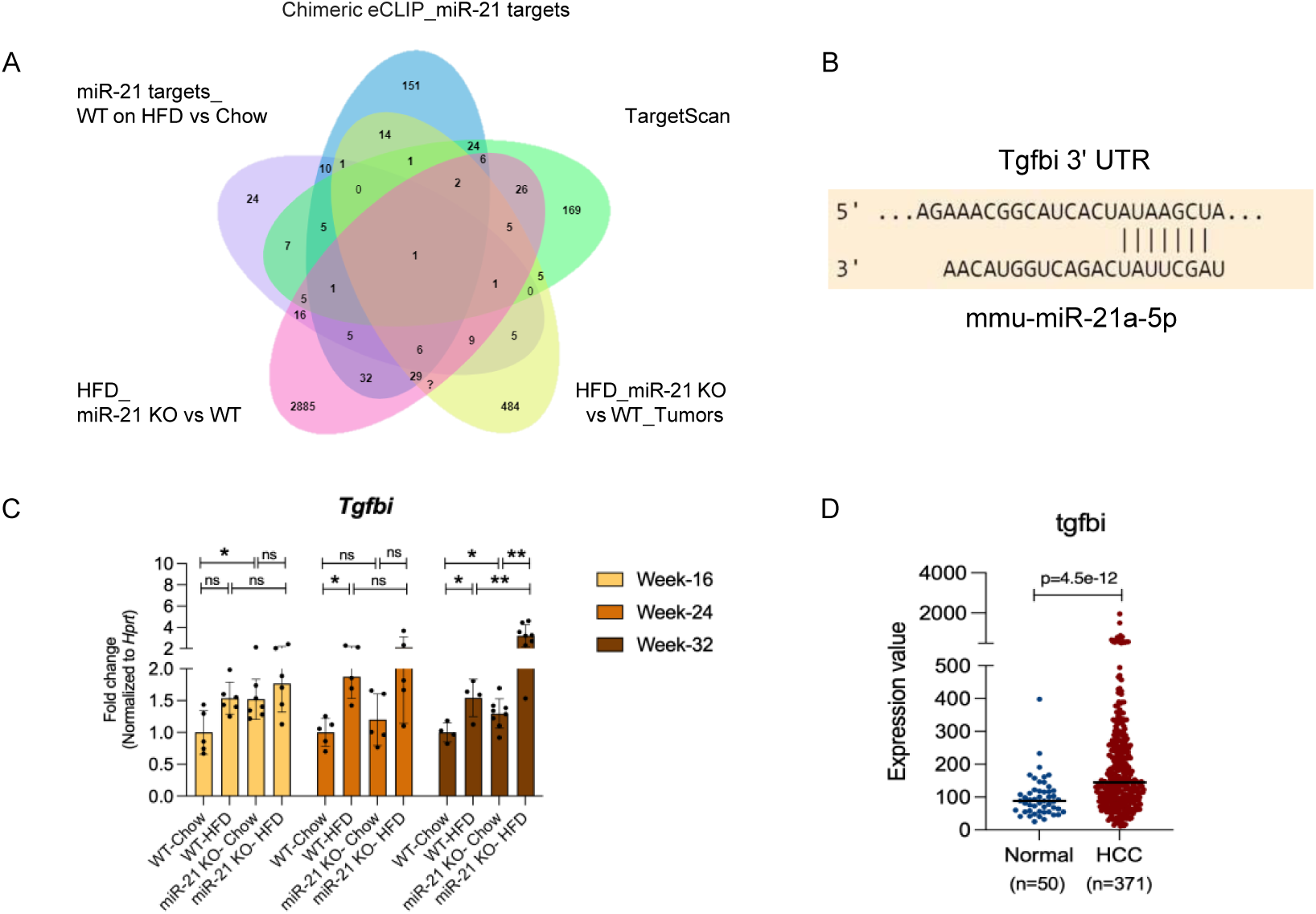
miR-21 directly regulates Tgfbi. (A) Venn diagram showing overlap of differentially expressed genes in various comparative datasets. (B) The in-silico binding of miR-21 on 3’UTR of *Tgfbi* gene, (C) Quantitative real-time analysis of *Tgfbi* mRNA expression in week 16, 24, and 32 liver samples. (D) *TGFBI* gene expression levels in the livers of HCC patients. Data is expressed as at least 4 mice/group. ns=not significant, *p<0.05, **p<0.01, ***p<0.001.

To identify direct targets of miR-21-5p in mouse liver, we performed chimeric eCLIP on liver tissue isolated from 8-week-old C57BL/6J mice, followed by probe capture for miR-21-5p, as described in the preprint by Manakov et al. (39). This analysis from an average of 3 replicates identified 484776 chimeric reads corresponding to 8618 genes (Suppl table 3). We compared this list to the sets of genes we found dysregulated in miR-21 KO livers, WT-HFD livers, hepatic tumors in HFD-fed mice, and the set of predicted miR-21 targets by TargetScan, 8.0 (40) and found substantial overlap in both cases (41)(Fig. 5A). The direct target of miR-21 commonly identified in this analysis was *Transforming growth factor-β-induced (Tgfbi)* (Fig. 5A). *Tgfbi* encodes an extracellular matrix protein that plays a role in various physiological and pathological conditions, including diabetes, corneal dystrophy and tumorigenesis (42). As predicted by TargetScan (Fig. 5B), we also observed enrichment in the miR-21 peaks on the *Tgfbi* 3’UTR (mm10) in our chimeric eCLIP experiment (Suppl Fig. 7A) that aligns perfectly with the miR-21 reverse seed sequence (Suppl. Fig. 7B). We observed that *Tgfbi* levels increased in our WT mouse livers as the disease progressed. Reinforcing the inverse relationship, miR-21 KO mice showed significantly higher expression of *Tgfbi* mRNA levels at all stages when compared to the WT-Chow counterparts (Fig. 5C). The miR-21 KO-HFD group showed the highest *Tgfbi* levels in week-32 samples, correlating its expression with the most severe liver disease phenotypes. We extracted gene expression data from OncoDB (43) for human patients diagnosed with hepatocellular carcinoma and as expected, found *TGFBI* levels to be significantly upregulated (Fig. 5D).

### Administration of miR-21 mimic slows the progression of liver disease pathogenesis in WT mice on HFD

As our previous results suggested an inverse relation between disease pathogenesis and miR-21 mRNA levels in WT mice, we asked whether supplementation of miR-21 after the manifestation of the disease would slow down disease pathogenesis in WT mice on HFD. To address this, we injected WT mice with 75mg/kg of DEN intraperitoneally at three weeks of age and continually fed them on HFD as before. This 3-fold increase in the amount of DEN, as compared to previous experiments, was used to accelerate the liver pathogenesis in an attempt to shorten the experimental timeframe. Based on the prior experiments, we allowed the mice to develop until week 16 and assumed that metabolic defects like obesity, insulin resistance, steatosis, and fibrosis had developed during this time, as seen before. At week 16, we randomly divided the mice into two groups and injected lipid nanoparticles encapsulating either the negative control (NC) or miR-21 mimic intravenously via tail vein injection. Each group was injected twice per week to administer a total of 1.8mg/kg of mimic each week for six weeks. At week 22, mice were sacrificed, and liver tissue and serum were collected for further analysis (Fig. 6A). As *Tgfbi* is a direct target of miR-21, we checked its levels as a response to miR-21 mimic administration. As expected, we observed a significant reduction in its mRNA levels as compared to the NC mimic (Fig. 6B).

**Figure 6:**
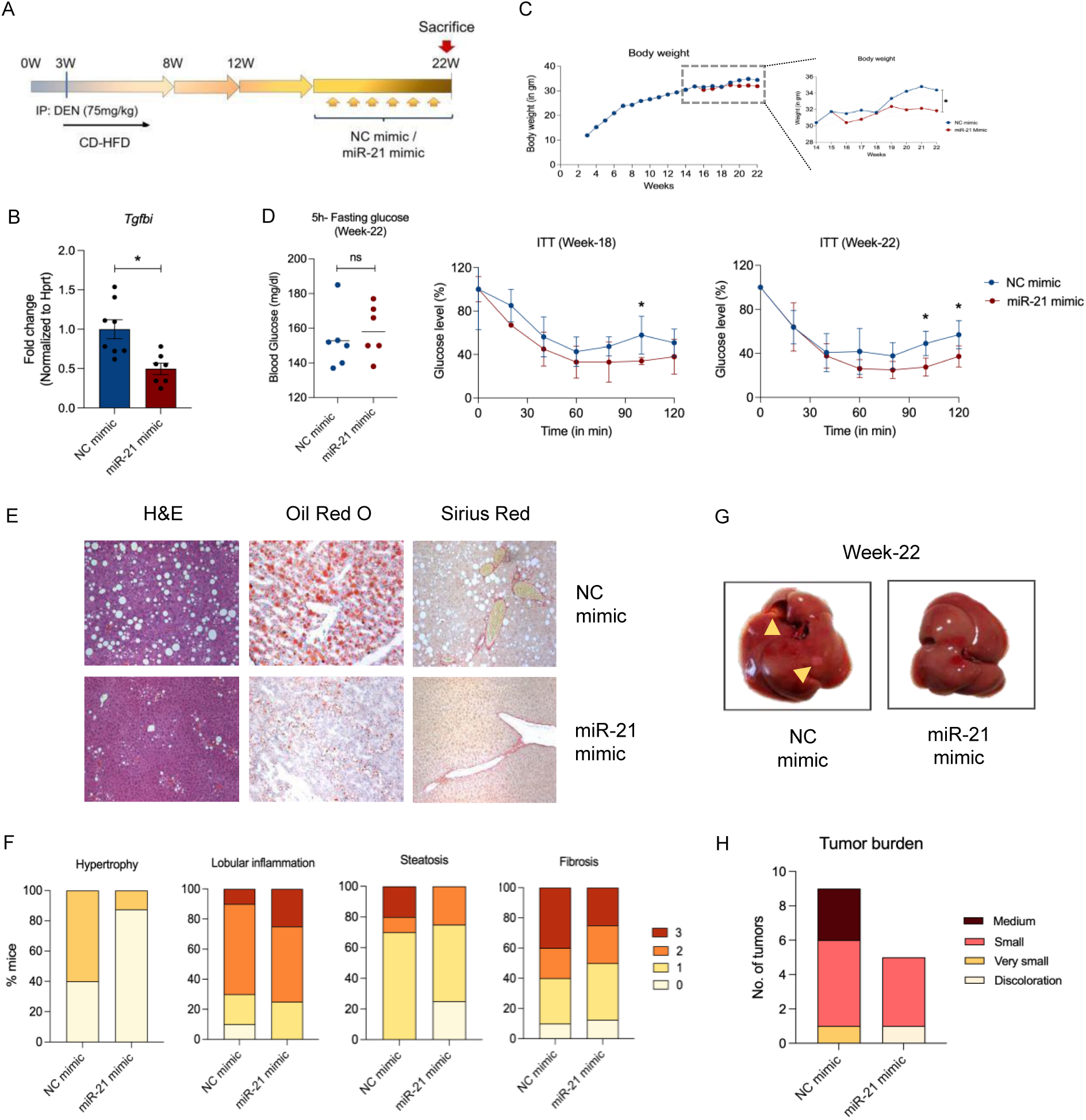
Supplementation of miR-21 mimic slows down the progression of liver disease in WT mice. (A) Timeline of the experiment (B) Quantitative real-time analysis for the levels of miR-21 target gene- *Tgfbi* in the liver (C) Body weight over time (D) Glucose levels in mices’ serum after 5-hours of starvation, Insulin tolerance test for week-18 and 22 (E) Histology of liver sections and staining with H&E, Oil Red O and Sirius red stain, (F) Histology scoring of Hypertrophy, Lobular inflammation, Steatosis and Fibrosis (G) Representative images of negative control (NC) and miR-21 mimic administered livers and (H) Tumor burden on week-22. ns= not significant. \**p*<0.05.

We then analyzed the metabolic phenotypes of these mice. Bodyweight analysis showed that the miR-21 mimic treated mice showed significant stability from further weight gain over the 6-week window (Fig. 6C). We further performed an insulin tolerance assay, which showed comparable glucose levels in the serum of both groups after 5 hours of starvation. However, upon insulin administration, the miR-21 mimic-injected mice showed a prolonged response to the presence of insulin as compared to the NC mimic group, indicating heightened insulin sensitivity at both week 18 and week 22 timepoints (Fig. 6D).

Further, we extracted the livers and performed H&E, Oil Red O, and Sirius red staining on liver sections to examine the overall liver architecture, steatosis, and fibrosis. Interestingly, the livers of miR-21 mimic treated mice showed healthier tissue architecture, reduced hypertrophy, reduced fat accumulation, and fibrotic tissue in their livers (Fig. 6E). The blinded histology scoring of these samples also supported these results, except for miR-21 mimic injected mice that showed slightly higher inflammation (Fig. 6F). Finally, we measured the tumor burden in these mice and observed that the miR-21 mimic injected mice showed reduced tumor burden as compared to the NC mimic group (Fig. 6G, 6H).

Together, these results indicate that administration of miR-21 as a mimic reduced the HFD-indued pathological features of liver disease. These results demonstrate a protective function of miR-21 in liver disease pathogenesis and point to miR-21 as a potential target to prevent/slow liver disease progression induced by high-fat exposure settings.

## Discussion

In this study, we have created a comprehensive, high-fat diet-based model of liver disease in mice that recapitulates the clinical stages observed in human patients, ranging from metabolic syndrome to MASLD to HCC. We have also analyzed the role of miR-21 in this disease progression. Among the most compelling findings of our study is the heightened susceptibility of miR-21 knockout mice to systemic insults, such as exposure to a low dose of DEN and a chronic high-fat diet (HFD). The accelerated disease progression and the increased severity of all metabolic and pathological stages of liver disease in the miR-21 KO-HFD combination reveal the protective role of miR-21 in liver disease progression *in vivo* under these metabolic conditions. In addition, our studies suggest that this protection by miR-21 is mediated through its direct gene target, *Tgfbi*. For the purpose of creating a liver disease model, we have used mice from the B6/129F2J background. It is well known that different strains exhibit substantial variations in MASH pathology and liver fibrosis. For example, analysis of clinical and molecular traits in 100 unique inbred mouse strains showed a >30-fold variation in hepatic TG accumulation (44) or various degrees of NAFLD (45). Recent studies analyzing different genetic backgrounds fed on varying dietary regimes, i.e., A/J, BALB/c, C3H/HeJ, C57BL/6J, CBA/CaH, DBA/2J, FVB/N and NOD/ShiLtJ (46) and C57BL/6J, DBA/2J, A/J, 129S1/SvlmJ, WSB/EiJ, CAST/EiJ, and PWK/PhJ (46) pointed towards the FVB/N mouse strain as the most adequate diet-induced mouse model for the recapitulation of MASLD (PMID: 36949095), and the PWK/PhJ mice as exhibiting the highest overlap with the human NASH signatures (46). Reportedly, diet-induced non-alcoholic/alcoholic fatty liver disease is most consistently promoted in 129Sv mice compared to C57BL/6 and CD-1 mice in terms of hepatic lipid profile (47) and in the diet-induced animal model of non-alcoholic fatty liver disease (DIAMOND), B6/129, a cross between two mouse strains, 129S1/SvImJ and C57Bl/6J, recapitulated non-alcoholic steatohepatitis (NASH) and hepatic cancers signatures, while the parent strains did not. While all the mentioned studies report that the extent of phenotypic changes triggered by environmental factors vary greatly based on the genetic background of the animal model, none of the studies include B6/129F2J, to the best of our understanding, a strain related to B6/129 that we used in this study.

It is well-established that a high-fat diet causes obesity and hepatic steatosis. Various HFD-based mouse models of liver disease exist (48–50); however, the lack of a complete spectrum of human phenotypes in these model organisms has hindered their acceptance. In addition, most of the proposed models require a long time to develop liver disease symptoms, especially severe liver disease pathologies like hepatocellular cancer. On the other hand, the models proposed to study the later stages either utilize high doses of carcinogens or genetic mutations. In this study, we attempted to create a model that would constitute the whole liver disease spectrum as well as an obesogenic diet that represents US dietary habits. To mimic the human obesogenic diet, we adopted a diet consisting of 58% kcals originating from fat. To accelerate the development of MASH-associated pathologies such as HCC while also minimizing the impact of the carcinogen, we limited the dose of DEN to 25mg/kg, using only a one-time injection.

In support of reports showing a protective role for miR-21 in HCC (34,35), our study provides additional evidence that challenge the conventional oncogenic role of miR-21. By utilizing the whole-body miR-21 knockout mouse model, we found that the loss of miR-21 leads to exacerbated liver injury, hepatomegaly, hyperglycemia, insulin resistance, steatosis, fibrosis, and, ultimately, HCC in mice. Our results unambiguously affirm the protective role of miR-21 during liver disease progression.

While we were able to demonstrate all features consistent with the various spectrum of metabolic phenotypes, i.e., obesity, hepatomegaly, hyperglycemia, and insulin resistance, as well as pathophysiological phenotypes, i.e., steatosis, fibrosis, MASH, and progressive hepatocellular carcinoma by high-fat-diet, we also showed that these effects were much more severe in the miR-21 KO background. This is consistent with the multiple-hit hypothesis of liver pathogenesis, where the loss of miR-21 may serve as the initial hit, rendering the liver more vulnerable to subsequent ‘secondary hits’ in the form of DEN, high-fat diet, and ensuing metabolic dysfunctions. Our study thus offers insights into the multifaceted interplay between genetic predisposition and environmental factors in liver disease progression to HCC.

Exploring the mechanisms underlying liver vulnerability in this multiple-hit model, we observed an inverse relationship between miR-21 levels and liver disease severity. Notably, the direct miR-21 target, *Tgfbi*, exhibited increased expression in the absence of miR-21, both in the presence and absence of HFD exposure. Previous literature has linked elevated *Tgfbi* levels in the serum of patients diagnosed with various gastrointestinal tract carcinomas (51), which, interestingly, showed reduced levels in patient cohorts that underwent treatment (52), emphasizing its role in cancer pathogenesis. In mouse models, *Tgfbi* KO mice showed resistance to adipose tissue hypertrophy, liver steatosis, and insulin resistance (53). In a separate study, the overexpression of *Tgfbi* showed an increase in the incidence of carcinogen-induced liver pre-neoplasia and tumors, whereas the knockout showed protection (52), demonstrating that it is causal in tumor formation. In our study, as expected from a direct miR-21 target gene, liver *Tgfbi* showed significant upregulation in response to miR-21 KO with and without HFD exposure. In fact, at week 32, while the tumor incidence was highest, the miR-21-HFD group showed even higher *Tgfbi* levels than all other groups. In contrast, *Tgfbi* levels were repressed by exogenous miR-21 administration. We further tested this correlation in HCC patients’ liver tissues and found it to be consistent with our data and previous literature (43). Our results strongly suggest that *Tgfbi* may be a pivotal downstream target that is responsible for promoting liver disease in the context of miR-21 loss, with or without concurrent HFD exposure. Coupled with previous literature showing its role in liver disease and cancer, this study makes *Tgfbi* a potential causal miR-21 downstream target promoting liver disease in the HFD setting.

MicroRNA-based therapeutics hold considerable promise as a strategy for restoring depleted microRNA levels. Owing to the observed reduced levels of miR-21 under severe liver disease conditions, we speculated that augmenting these microRNA levels might have a beneficial impact on disease progression. Our hypothesis was also supported bya study by Said Lhamyani et al., where the authors demonstrated that the miR-21 mimic administration delayed weight gain in HFD-fed mice compared to the control littermates, placing miR-21 mimic as a potential therapeutic tool to modulate adipose function and control obesity (35). To investigate our hypothesis, we employed lipid nanoparticles to encapsulate the miR-21 mimic and administered them by tail vein injections to wild-type mice that had already manifested symptoms of metabolic syndrome, steatosis, and fibrosis. Our weekly six-week injection regimen demonstrated a significant improvement in various parameters, including body weight, insulin sensitivity and levels of steatosis and fibrosis. Our study underscores the potential of miR-21 as a viable microRNA therapeutic target for the prevention and treatment of liver disease.

Clinical significance – We propose that the potential mechanism of toxicity of miR-21 KO and HFD combination is through increasing the levels of miR-21 target protein, *Tgfbi*. The clinical samples of HCC tissues showed a similar significant increase in the levels of this gene correlating with HCC phenotypes, as observed in our results. We propose that similar to targeting miR-21, targeting these genes could also be a potential mechanism to tackle liver disease progression. Our study has important implications for understanding the mechanisms underlying liver disease progression, especially in patients with low expression of miR-21 and a lifestyle with fatty, obesogenic diets, and for the design of future studies directed at the prevention and treatment of liver disease.

Limitations of the study-A limitation of this study is that the knockout of miR-21 is in the whole body of mice, and therefore, the effects caused specifically by the knockout of miR-21 in the liver, or a specific liver cell type were not determined in our study. However, previous work has revealed that miR-21 KO in just hepatocytes was associated with increase liver cancer in a tissue-specific KO mouse model (34). We note that the mechanism of action of miR-21 to protect from liver disease requires additional experimentation that addresses the cause-and-effect relationship between the various players involved.

Taken together, our study introduces a comprehensive murine model of liver disease based on a high-fat diet and a small amount of carcinogen, encompassing a spectrum of human phenotypes and highlighting the protective role of miR-21. Using whole-body knockout of miR-21, we showed that the animals are predisposed to upcoming liver insults, and this loss is sufficient to cause notable liver disease pathogenesis. In addition, we show that the combination of miR-21 KO and HFD is a recipe for severely accelerated liver progression from simple steatosis to hepatocellular carcinoma, thus highlighting the protective role of miR-21 in prevailing Western diet metabolic settings. We further suggest that this protection by miR-21 is mediated, at least in part, by regulating its direct target, *Tgfbi*. Finally, we propose that targeting miR-21 as a therapeutic might be an effective strategy to prevent or delay liver disease, including HCC.

While acknowledging the limitations of our study, we anticipate that our findings will stimulate further investigation into intricate mechanisms governing liver disease and pave the way for innovative therapeutic strategies. Our study has far-reaching implications, not only in advancing the understanding of liver disease pathogenesis but also in guiding future research and therapeutic interventions aimed at preventing and treating these debilitating conditions.

## Materials and methods

### Sex as a biological variable

Our study examined male mice as male animals are more likely to be affected by severe liver disease.

### Animals

The B6/129SF2/J mice were purchased from Jackson Laboratory and were used as wild-type (WT) controls. miR-21 KO (B6;129S6-*Mir21atm1Yoli*/J) mice with the same background were used for the experiment. All mice were housed in a 12h light – 12h dark cycle in a 21-23°C pathogen-free barrier facility with free access to water. All procedureswere performed according to protocols approved by the Animal Care facility of BIDMC and IACUC (Protocol number: 010-2021).

### Dietary interventions

Male mice were used for this study. At three weeks of age, all mice were injected with 25mg/kg of Diethylnitrosamine (DEN) (Cat #N0756) by intraperitoneal injections. All mice were maintained on *ad libitum* of either a normal chow diet (Research Diets, D12331, %kcal: 25.7% Protein, 62.9% Carbohydrate, 11.4% Fat) or high-fat diet, without additional choline(Research Diets, D17022005i, %kcal:17% Protein, 26% Carbohydrate, 58% Fat). Full composition: Casein, 80 Mesh, 228gm; DL-Methionine, 2gm, Maltodextrin 10, 170gm; Sucrose, 175gm, Soyabean Oil, 25gm; Coconut Oil, Hydrogenated, 333.5gm; Mineral Mix S10001, 40gm; Sodium Bicarbonate, 10.5gm; Potassium Citrate, 1 H2O, 4gm; Vitamin Mix V10001; 10gm; FD&C Red Dye #40; 0.1gm. The mice were euthanized at their respective time points for further analysis.

### Biochemical analysis

Blood was collected at regular intervals for biochemical analysis using the submandibular blood collection method. Glucose measurement was done using the Accu-Chek Guide. ALT (Teco diagnostics, Anaheim, USA), AST (843 3815), Triglyceride (133 6544), and cholesterol (166 9829) measurements were done using the instrument VITROS350 from Ortho-Clinical Diagnostics. Whole body fat was measured using the EchoMRI^TM^ 2009 3-in-1 Composition analyzer (Version 130718).

### Insulin tolerance assay

All the mice were fasted for 5 hours before performing the Insulin tolerance assay (ITT). The mice were injected with an insulin solution (1.5U/kg) via intraperitoneal injections of Humulin R – NDC 0002-8215-17. A minimum of 5 mice per group was used for the assay. During the assay, mice were measured for glucose at 0 min (baseline), 15 min, 30 min, 45 min, 60 min, 75 min, 90 min, 105 min, and 120 minutes post-insulin administration by tail-snip. Insulin resistance (IR) was determined using the homeostasis model assessment for insulin resistance (HOMA-IR; (fasting insulin* fasting glucose)/22.5).

### Histological analysis

For liver histological analysis, mouse liver tissues from weeks 16, 24 and 32 timepoints were dissected and fixed in either 10% neutral buffered formalin and subsequently embedded in paraffin or placed in OCT and frozen on dry ice. Liver histology was assessed using Hematoxylin and Eosin (H&E) staining in paraffin-embedded sections at the BIDMC institutional core facility. Fibrosis was assessed using Sirius red staining in paraffin-embedded sections. The presence of steatosis was confirmed with Oil-Red-O stains in frozen sections. An expert pathologist evaluated the liver histology in a blinded manner. The Histology Scoring systems from Liang et al. (54) and Kleiner et al. (55) were adapted to assess the phenotypes. The liver sections were evaluated for the presence and degree of steatosis, inflammation and fibrosis. The macroscopic and microscopic tumor nodules were elevated and classified according to their malignant potential. A panel of immunohistochemical (IHC) stains used to study malignant potential included glypican 3 (Invitrogen PA5-89561, polyclonal, 1:300 and 1:600 dilutions), glutamine synthetase 6 (Abcam AB176562, clone EPR13022, 1:500 dilution), β-catenin (Abcam AB223075, clone IGX4794R-3, 1:4000 dilution), heat shock protein 70 (Abcam AB194360, clone EPR16893, 1:4000 dilution), and serum amyloid A (Bio SB BSB2806, clone EP335, 1:100 dilution). All IHC staining was performed on the Leica Bond III platform.

### RNA extraction, cDNA preparation, and quantitative RT-PCR

Total RNA extraction and quantitative real-time PCR (qRT-PCR) were performed using Trizol (Cat #15596018, Invitrogen). The primers were purchased from Integrated DNA Technologies, USA. Primer sequences are provided in the supporting table 1. For miR-21 quantification, the pre-designed miRCURY LNA miRNA PCR assay kit from Qiagen was used (GeneGlobe ID - YP00204230, Catalog No. 339306).

### Library preparation and sequencing

Libraries were prepared from the extracted RNA using the KAPA RNA HyperPrep Kit with RiboErase (HMR)- KK8561. To understand the effects caused by HFD exposure, and miR-21 KO independently and in combination, we performed a bulk RNAseq analysis. Livers from all four groups, i.e., WT-Chow, WT-HFD, miR-21 KO-Chow, and miR-21 KO-HFD at weeks 16, 24, and 32, were extracted and were used to isolate RNA. In addition, tumors extracted from WT-HFD and miR-21 KO-HFD at week 32 were also used to isolate RNA and perform bulk RNA sequencing.

### RNA-seq analysis

Raw sequencing reads were aligned to a reference transcriptome generated from the Ensembl v104 GRCm38 database with salmon v1.4.0 using options “–seqBias – useVBOpt –gcBias –posBias –numBootstraps 30 –validate Mappings”. Length-scaled transcripts per million were acquired using tximport v1.18.0, and log2 fold changes and false discovery rates were determined by DESeq2 in R. T-distributed stochastic neighbor embedding (Tsne) was performed with counts transformed by the variance Stabilizing Transformation function from DESeq2.

### Differential expression analysis

Qiagen Ingenuity Pathway Analysis (IPA)-94302991 software was used to perform gene enrichment and comparative analyses for all samples. A part of the analysis was also performed using the online tool Database for Annotation, Visualization, and Integrated Discovery (DAVID) software.

### Lipid nanoparticle formulation

mirVana mimics purchased from ThermoFisher were used in the synthesis of miRNA-LNP. Mimics for miR-21a-5p (5’-UAGCUUAUCAGACUGAUGUUGA-3’) (MC10206), and Negative Control #1 (CM00102) were generated with a 5’-FAM modification. The miRNA mimics were dissolved in PNI formulation buffer (NWW0043) at a 0.24 mg/ml working concentration for an N/P ratio of 3. Encapsulation was performed using the NanoAssemblr Benchtop system (Precision NanoSystems). Diluted RNA was mixed with Genvoy-ILM-DiD (NWW0039), a fluorescently labelled, off-the-shelf lipid mix (molar ratios of 50:10:37.5:2.5 for PNI-Lipid, DSPC, cholesterol, and PNI Stabilizer). Components were mixed via a microfluidic cartridge (NIT0062) at a flow rate ratio of 3:1 (RNA in PNI buffer: Genvoy-ILM) at a total flow rate of 12 ml/min. After encapsulation, particles were dialyzed against PBS and concentrated using Amicon Ultra-15 Centrifugal Filters (EMD Millipore).

### Ribogreen Assay for RNA Encapsulation Efficiency

A modified Ribogreen assay (Invitrogen, Quant-iT Ribogreen RNA assay kit) was performed to determine the encapsulation efficiency. In a 96-well clear microplate, miRNA-LNP samples were added in a 1:1 ratio to either 1x TE buffer or 2% Triton buffer (Triton in TE Buffer) in duplicates with Ribogreen reagent. After a 10-minute incubation at 37 °C, the plate was read using a fluorescent plate reader with an excitation wavelength of 485nm and an emission wavelength of 528nm. Encapsulation efficiency and mRNA concentration were calculated using the standard curve and sample values.

### LNP Characterization

Mean diameter and polydispersity index (PDI) were determined by Dynamic Light Scattering (DLS) using ZetaSizer Nano ZS (Malvern Panalytical). Samples were diluted to ∼18ng/µl of total RNA and added to a quartz cuvette (ZEN0040), and measurements were made using a 173° backscatter angle. Measurement parameters were as follows: the material refractive index was 1.45m and absorption was 0.001; for the dispersant, viscosity was 1.0200 cP, RI 1.335, and dielectric constant 80.4. Ten runs of 10 seconds duration were performed for each of the measurements. Data was processed using a General-Purpose model with normal resolution.

### LNP injections in mice

For the therapeutic intervention experiment, the WT mice were injected with 75mg/kg of DEN at week three of age and were fed on HFD throughout the experimental duration. At week 16, the mice were divided into two categories: Negative Control (NC) and miR-21 mimic. Lipid nanoparticles encapsulating either the NC or miR-21 mimic were injected via the intravenous tail vein in each mouse twice weekly, totaling 1.8 mg/kg of mimic per week. The injections were performed for six weeks, and at the end of week 22, all mice were euthanized to collect livers.

### Statistical analysis

All data represent at least three independent experiments. P<0.05 was considered significant and indicated by *p<0.05, **p<0.01, ***p<0.001, ****p<0.0001. Analyses performed included two-way analysis of variance, Student’s *t*-test, and unpaired *t*-test where appropriate.

## Supplementary material

Figures – 7

Tables – 3

1. Primer sequences used for qRT-PCR analysis
2. Differential expression of genes in tumor samples from miR-21 KO vs WT on HFD and Cancermine analysis
3. miR-21 eCLIP target data

## Author contributions

FJS conceived, supervised, and secured funding for the project. UJ designed and performed experiments as well as wrote the manuscript. AQ assisted in all the in-vivo mouse experiments and led lipid nanoparticle formulations. JDL performed RNAseq mapping. YO and IN performed histology analysis and scoring. GS provided intellectual support throughout the study.

## Acknowledgements

We thank Drs Yuan Zhuang, Mrigya Babuta, and Prashant Thevkar for the discussion and critical input. We also thank Dr. John Clohessy and Dr. Maria Mavrikaki for their assistance during the experiments. We thank Teri Bowman, HT ASCP, who assisted with immunohistochemical staining for this project. The graphical abstract was created using BioRender.com. Graphs with appropriate statistical analyses were created using GraphPad Prism version 10.0.2 (171) for MacOS, GraphPad Software, Boston, Massachusetts, USA, www.graphpad.com.

## Abbreviations

ALT: Alanine aminotransferaseB6 B6/129SF2/J
DEN: Diethylnitrosamine
HFD: High-fat diet without additional choline
FDR: False discovery rate
ITT: Insulin tolerance test
MASLD: Metabolic dysfunction-associated steatotic liver disease (Formerly NAFLD)
MASH: Metabolic-associated steatohepatitis (Formerly NASH)
T2DM: Type 2 diabetes mellitus
HCC: Hepatocellular carcinoma

