## Supplementary figures for "miR-21: A therapeutic target for delaying severe liver disease and hepatocellular carcinoma in high-fat-diet-fed mice"

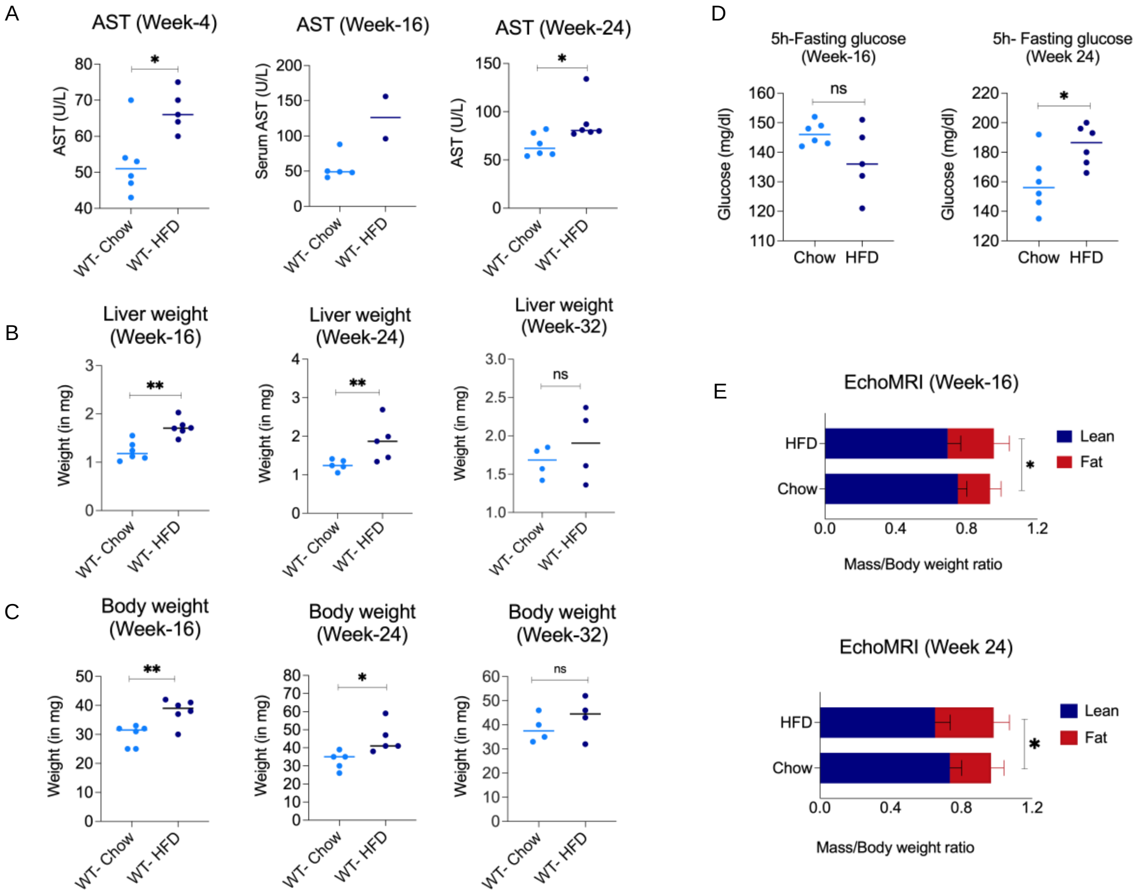


**Supplementary Figure 1: WT mice fed on HFD develop obesity, liver injury, hyperglycemia, and increased body fat content** (A) Serum levels of alanine transaminase (ALT), (B) Liver weight (C) Body weight at week-16, 24 and 32, (D) 5-hours of fasting blood glucose levels ratio (Normalized to WT-Chow group) before Insulin tolerance assay. (E) EchoMRI measurement for body fat composition Data is expressed as the mean±SD. ns=not significant, **p*<0.05, ***p*<0.01, ****p*<0.001.

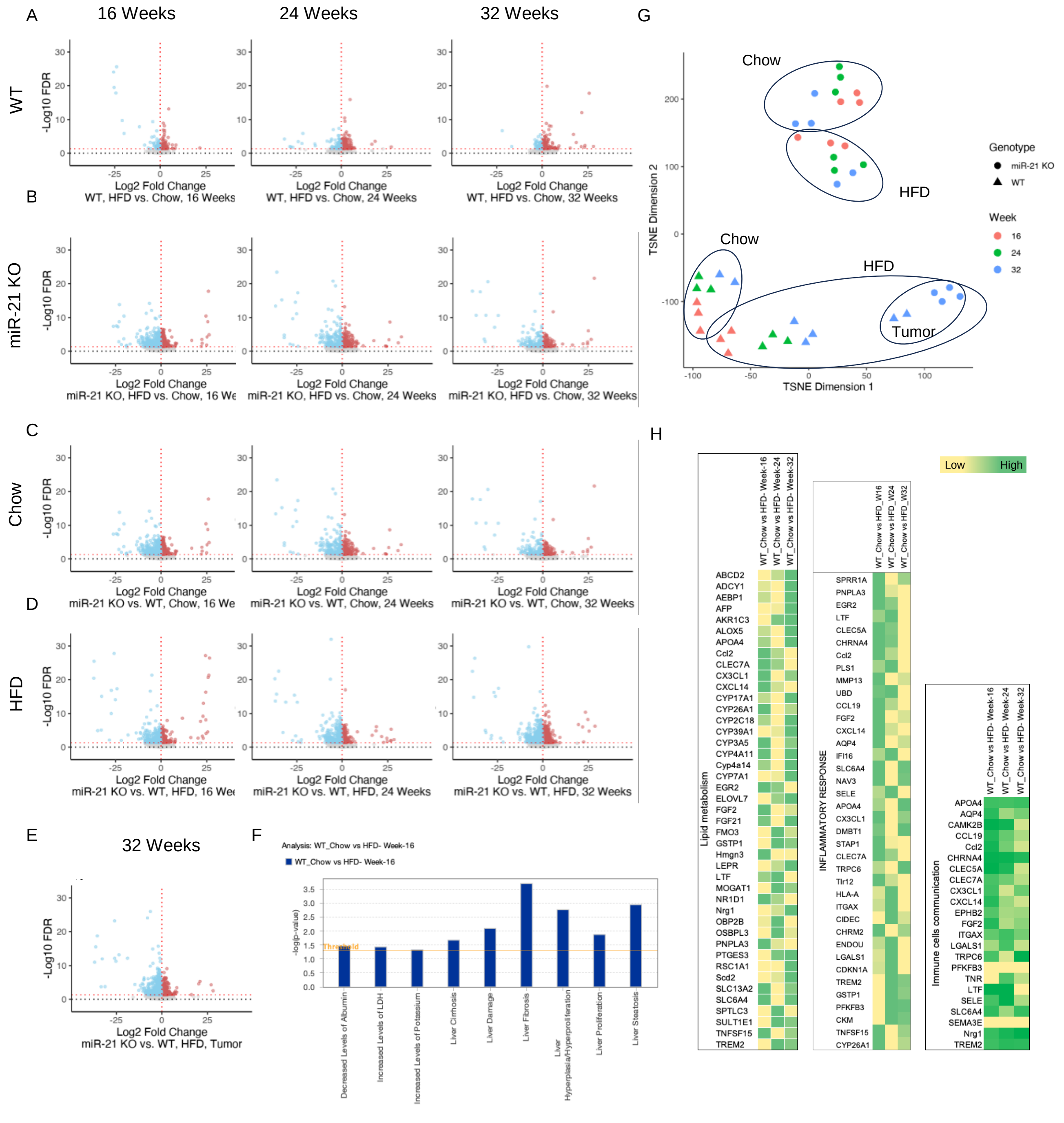


**Supplementary Figure 2: Transcriptomic analysis for RNAseq data on liver samples** -Log10 FDR values for differentially expressed genes with fold change > 2 at week-16, week-24, and week-32 for comparison of HFD vs Chow fed to (A) WT and (B) miR-21 mice, comparison of miR-21 KO vs WT mice fed on (C) Chow and (D) HFD, and (E) tumors isolated from miR-21 KO vs WT mice livers fed on HFD. (F) t-distributed Stochastic Neighbor Embedding (t-SNE) plot to visualize the RNAseq datasets. (G) Bar graph showing liver toxicity pathways being upregulated upon HFD exposure to WT mice (Threshold of –log(p-value)= 1.3), (H) Heatmap of differentially expressed genes involved in lipid metabolism, inflammatory response, and immune cell communication.

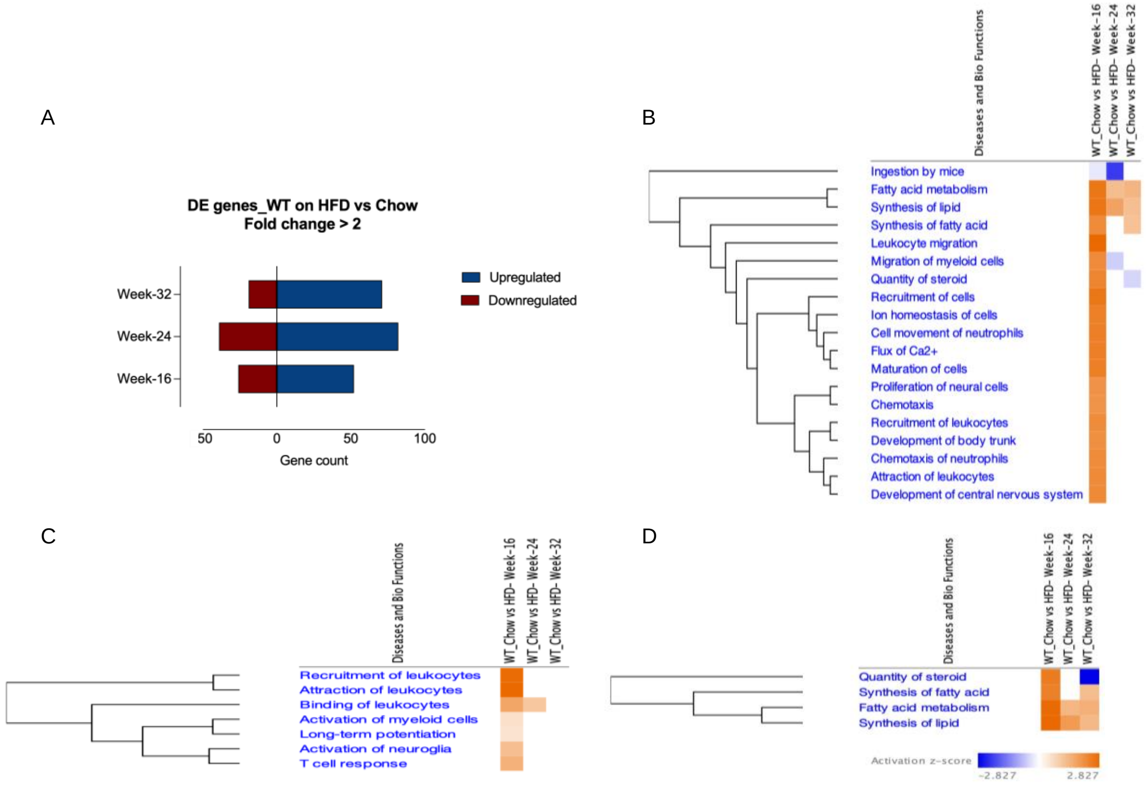


**Supplementary Figure 3: WT mice on HFD show a heightened metabolic response in the earliest time tested for hepatic injury, which stabilizes with time.** (A) Bar graph indicating differentially expressed up and down-regulated genes (fold change>2), (B, C, D) Activation Z scores for predicted activation or inhibition of various Diseases and biofunctions in WT livers upon HFD consumption at week 16, 24, and 32.

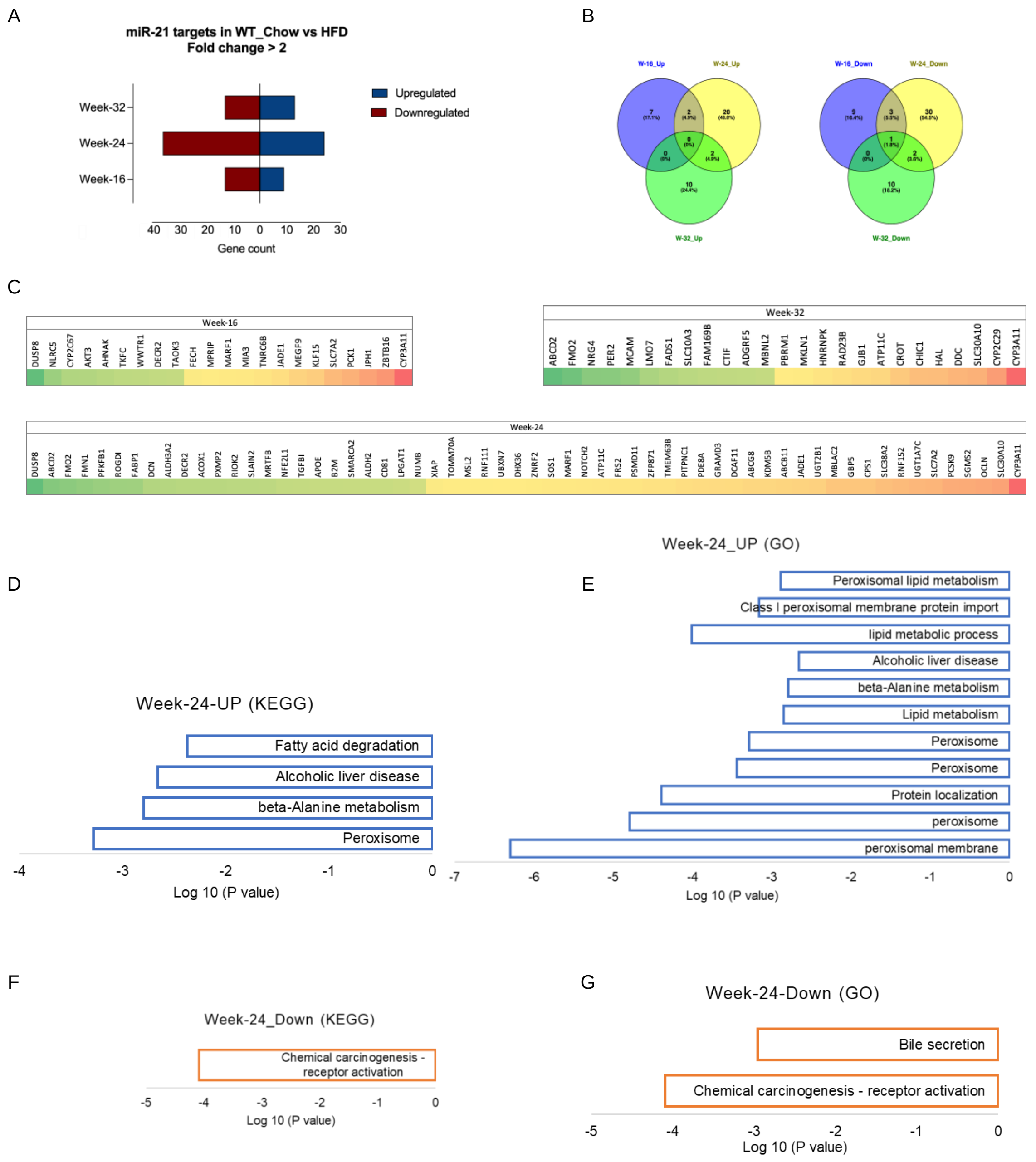


**Supplementary Figure 4:** Expression of miR-21 targets in WT mice upon HFD feeding (a) Gene counts of miR-21 targets in WT mice upon HFD exposure ($p<0.05$), (B) Venn diagram showing the overlap between different datasets, DAVID analysis showing Upregulated (D) KEGG and (E) GO pathways and downregulated (F) KEGG and (G) GO pathways in week-24 liver samples.

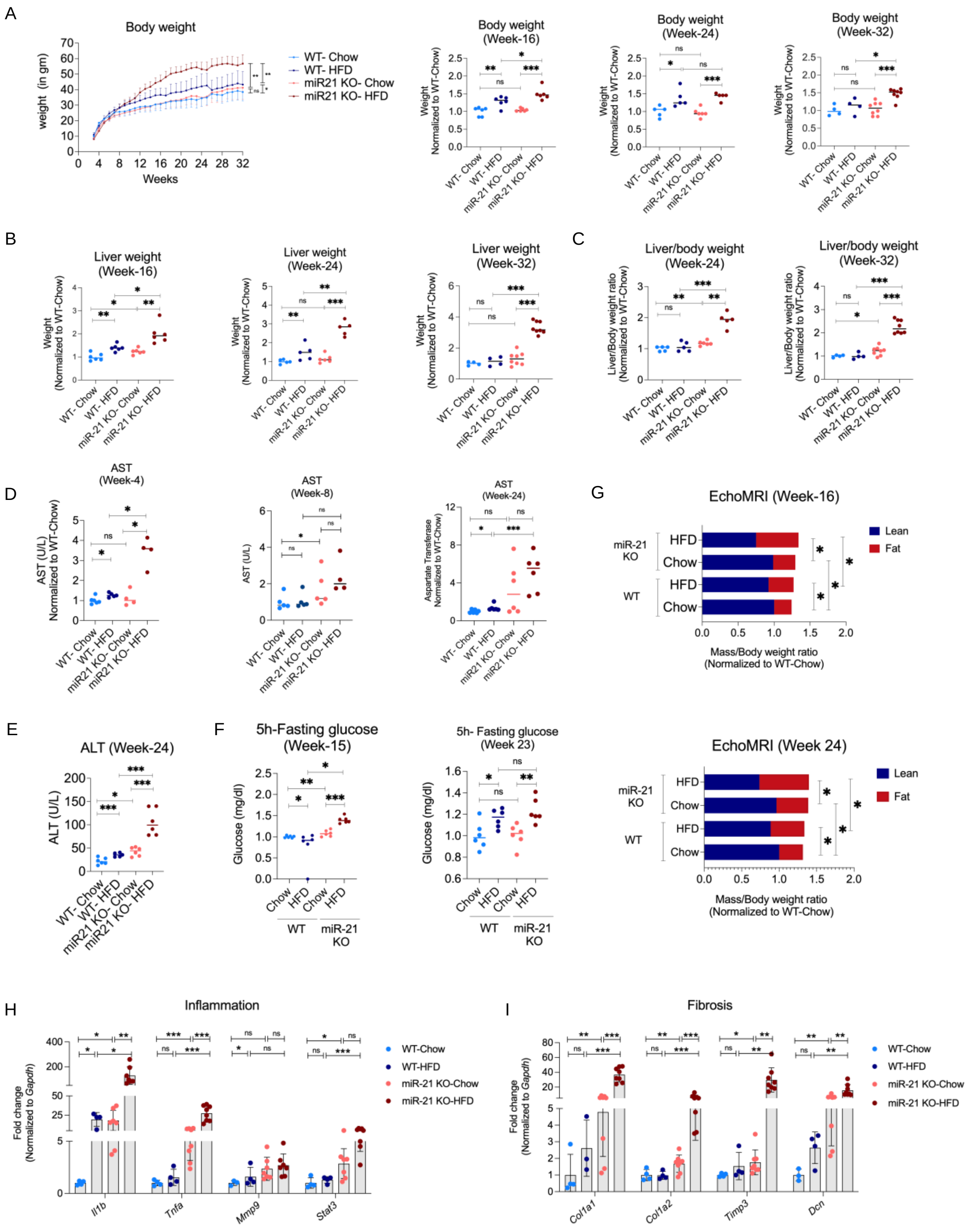


**Supplementary Figure 5: miR-21 KO mice and HFD exposure show additive effects on the development of obesity, liver injury, hyperglycemia, and increased body fat content** (A) Body weight measurement at all time points and normalized to WT-Chow at week-16, 24, and 32. (B) Liver weight at week-16, 24, and 32 (C) Liver to body weight ratio at week-24 and 32, (D) Serum levels of aspartate aminotransferase (AST) at week-4, and 24, and of (E) alanine transaminase (ALT) at week-24, (F) 5 hours of fasting blood glucose levels ratio as measured before Insulin tolerance assay. (G) EchoMRI measurement for body fat composition. Relative expression levels of genes involved in (H) inflammation and (I) fibrosis at week-32. All the data is normalized with the WT-Chow group and is expressed as the mean±SD. ns=not significant, *$p$<0.05, **$p$<0.01, ***$p$<0.001.

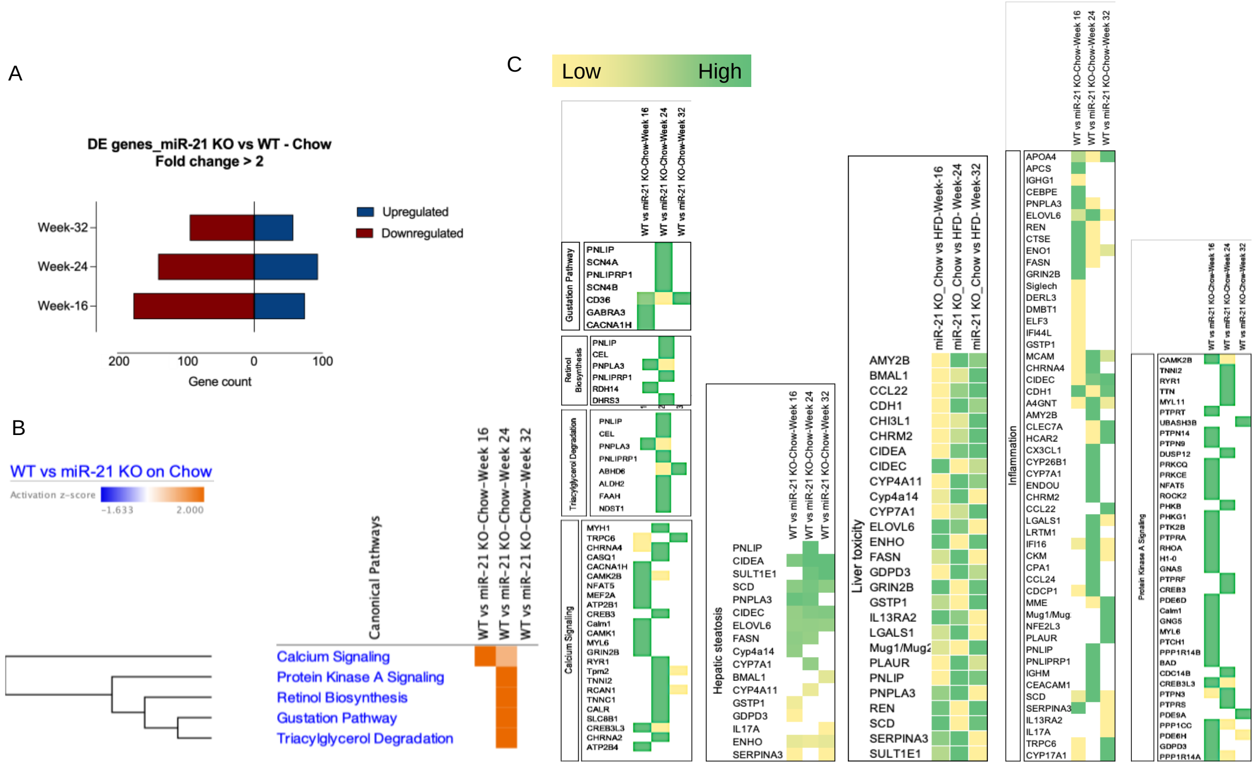


**Supplementary Figure 6: miR-21 KO mice are predisposed to metabolic syndrome and liver disease** (A) Gene count of differentially expressed genes with fold change > 2 in miR-21 KO mice as compared to WT fed on Chow, (B) IPA analysis of differentially expressed canonical pathways in miR-21 KO mice over time, (C) Heatmap representing the genes involved in calcium signaling, hepatic steatosis, liver toxicity, and inflammation Retinol Biosynthesis, Triacylglycerol degradation, Gustation pathway, and Protein Kinase A signaling.

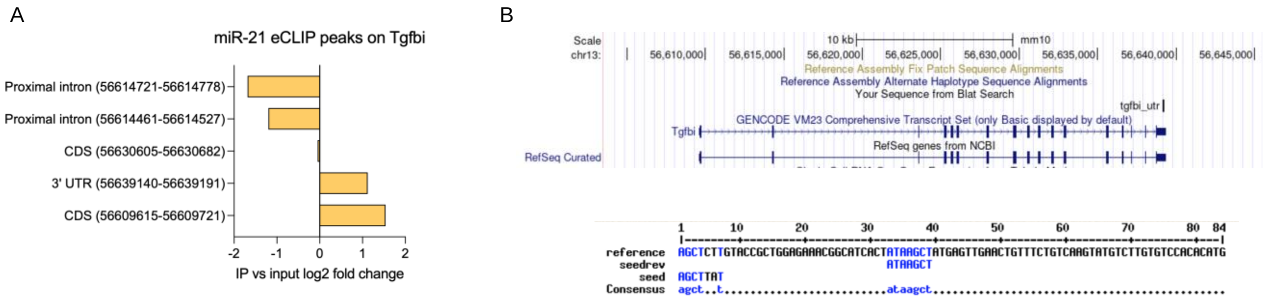


**Supplementary Figure 7: miR-21 binds directly to *Tgfbi*** (A) Log2 fold change values of miR-21 eCLIP peaks on *Tgfbi* genomic region, (B) Alignment of miR-21 seed sequence to the 3’UTR of *Tgfbi* gene (mm10) identified in the eCLIP read.
